# Beyond ingredients: Supramolecular structure of lipid droplets in infant formula affects metabolic and brain function in mouse models

**DOI:** 10.1101/2023.02.24.529900

**Authors:** Annemarie Oosting, Louise Harvey, Silvia Ringler, Gertjan van Dijk, Lidewij Schipper

## Abstract

Human milk beneficially affects infant growth and brain development. The supramolecular structure of lipid globules in human milk i.e., large lipid globules covered by the milk fat globule membrane, is believed to contribute to this effect, in addition to the supply of functional ingredients.

Three preclinical (mouse) experiments were performed to study the effects of infant formula mimicking the supramolecular structure of human milk lipid globules on brain and metabolic health outcomes. From postnatal day 16 to 42, mouse offspring were exposed to a diet containing infant formula with large, phospholipid-coated lipid droplets (structure, STR) or infant formula with the same ingredients but lacking the unique structural properties as observed in human milk (ingredient, ING). Subsequently, in Study 1, the fatty acid composition in liver and brain membranes was measured, and expression of hippocampal molecular markers were analyzed. In Study 2 and 3 adult (Western style diet-induced) body fat accumulation and cognitive function were evaluated.

Animals exposed to STR compared to ING showed improved omega-3 fatty acid accumulation in liver and brain, and higher expression of brain myelin-associated glycoprotein. Early exposure to STR reduced fat mass accumulation in adulthood; the effect was more pronounced in animals exposed to a Western style diet. Additionally, mice exposed to STR demonstrated better memory performance later in life.

In conclusion, early life exposure to infant formula containing large, phospholipid-coated lipid droplets, closer to the supramolecular structure of lipid globules in human milk, positively affects adult brain and metabolic health outcomes in pre-clinical animal models.

## Introduction

It is well established that breastfeeding brings many health benefits to both mother (1, 2) and infant. Acute benefits for infants include protection against neonatal morbidity and mortality (3) and against proinflammatory gastrointestinal conditions and infections (4, 5), while long-term effects include reduced incidence of asthma (6) and (food) allergies, improved cardio-metabolic health and reduced obesity risk (7–9), and improved neurocognitive functions (10, 11). Many aspects of breastfeeding may contribute to these health benefits. Aside from the difference in feeding mode (i.e., breast versus bottle feeding (12–14)) and nutritional composition tailored to age-specific and individual needs (15), human milk (HM) contains bioactive compounds that contribute to its functional benefits (15–17) over infant formula (IF). These compounds include hormones, growth factors, and immunomodulators, which can all contribute to healthy growth, gut maturation, microbiota, and development of the brain and immune system (15).

Selected bioactive compounds can and have been added to IF to bring the nutritional composition of IF closer to that of HM (16, 18). However, differences between IF and HM remain, including important differences in the dietary lipid fraction. Lipids are an important constituent of milk, providing energy and building blocks that can be used as structural components of cell membranes (19). In mammalian milk, it is a complex mixture of different types of lipids structured as milk fat globules (MFGs; Fig 1A). These are formed in the mammary epithelial cells and secreted through exocytosis in the alveolar lumen, resulting in globules with a core of triglycerides (TG) and cholesteryl esters (CE) (98% of total lipids) surrounded by a native biological membrane composed mainly of phospholipids, proteins and enzymes, free cholesterol, glycolipids and glycoproteins - the milk fat globule membrane (MFGM). The MFGM accounts for 2-6% of the total MFG mass, of which 25-60% consists of MFGM proteins (1-4% of total milk protein) and polar lipids contributing 0.2-1% to total milk lipid content (20, 21). The lipid globule size ranges between 1 and 10 μm with an average mode diameter of 4 μm in mature milk (22, 23).

**Figure 1.**
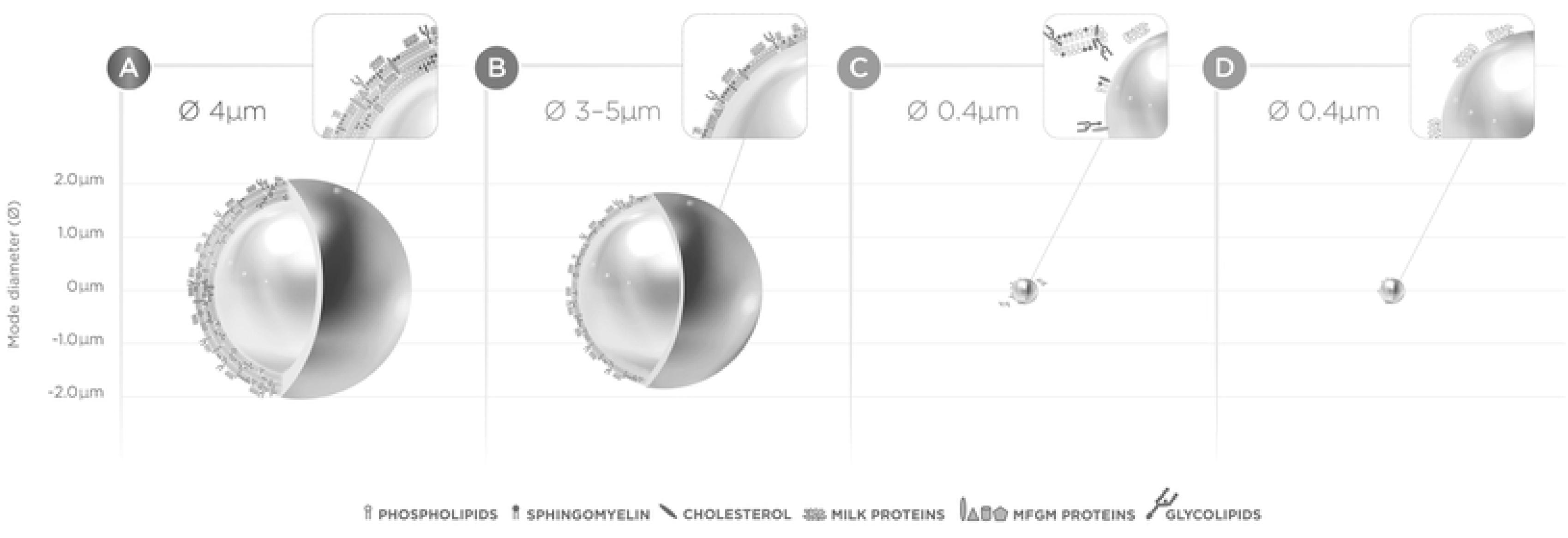
Schematic overview of lipid globule with tri-layer of MFGM in human milk (A), lipid droplet with MFGM fragments at the droplet interface (B; Nuturis), lipid droplet in standard infant formula with MGFM dry blended (C) and standard infant formula without MGFM but with milk proteins at the droplet interface (D). Visual representation of the size and ratios of different MFGM and milk protein molecules does not fully reflect reality.

In contrast to the complex supramolecular structure observed in HM, lipid droplets in IF have a much smaller mode diameter of approximately 0.4 μm and consist of a triglyceride core with casein and whey milk protein adhering to the globule surface ((22, 24, 25); Fig 1D). IF may contain low concentrations of vegetable lecithin added as an anti-oxidant and/or as an emulsifier, but contains no or only traces of sphingomyelin nor cholesterol, unless dairy lipids are used as ingredients (22). Although the primary function of MFGM is to allow the TG core to be secreted in the alveolar lumen and remain in a stable emulsion in the milk, it has been suggested that its constituents are involved in many biological functions, including cell signalling and growth, anti-microbial and anti-viral defence, apoptosis, differentiation, myelination and energy metabolism (20, 21). For this reason, adding MFGM fragments as an ingredient to IF could potentially contribute to improved neurodevelopment, gut maturation, metabolism and immunity of formula fed (FF) infants, bringing their developmental trajectory closer to that of BF infants (26, 27). Indeed, a recent meta-analysis by Ambrozej et al (28) showed, despite limited clinical evidence, that MFGM-supplemented formula reduced otitis media infection incidence and improved cognitive development compared to non-supplemented formula. These reported clinical benefits suggest that adding MFGM narrows the gap between HM and IF composition and function.

However, it is important to highlight that there is more to the structure of human MFG than only MFGM. While solely adding MFGM fragments to IF provides functional membrane lipids and proteins, it does *not* result in the same MFG *structure* as found in HM. Isolating dairy MFGM from bovine milk disrupts the membrane and results in ingredients enriched in MFGM fragments (29, 30). Subsequent additions of a MFGM ingredient to standard IF will result in small lipid droplets, with the MFGM fragments mainly remaining in the aqueous phase, and only limited amounts of phospholipids at the interface of the lipid droplets ((31, 32), Fig 1B). It is thought that both globule size and the MFGM at the interface of the MFG affect lipid digestion and absorption kinetics, proteolysis, plasma TG appearance and β-oxidation (33–35). Given the dependence of organs and tissues in early life on dietary lipids as energy and building blocks, differential bioavailability of these components after ingestion may impact an infant’s developmental trajectory and (long-term) function. Thus, the supramolecular structure of lipid droplets in infant nutrition is hypothesized to affect infant metabolic and neurocognitive development and (long-term) function, possibly contributing to some of the developmental differences observed between BF and FF infants.

A concept IF (Nuturis) was developed to narrow the gap between standard IF and human MFG structure addressing both the size and the interfacial composition. It more closely mimics the human MFG by presenting lipid droplets with associated HM lipid components (such as phospholipids, cholesterol and MFGM fragments) and a volume-weighted mode diameter of 3-5 mm in line with that of human milk, despite not providing a true tri-layer MFGM ((22, 24, 25); Fig 1C). To establish the distinct effect of early life exposure to supramolecular lipid structure on brain and metabolic health outcomes, while excluding the effect of added phospholipids, we performed three mouse experiments using either egg or milk derived phospholipids.

## Materials and methods

Three separate studies were conducted, using mouse models with a roughly similar design, at three different laboratories in the Netherlands: All studies included an exposure to experimental diets varying in lipid droplet structure during a standardized period early in life, but other experimental conditions (e.g. diets, husbandry), experimental designs, and the readout parameters, differed per study, see below. Study 1 aimed to investigate the impact of lipid droplet structure on bioavailability of fatty acids (FA) and their incorporation in functional membranes. Study 2 and 3 investigated the long-term effects of early life exposure to lipid droplet structure on adult metabolic health and cognitive function, respectively.

### Animals & care

All animal procedures were performed according to general guidelines for the care and use of experimental animals, and the experiments were compliant to the EU Directive on the protection of animals used for scientific purposes and the Dutch Animal Welfare Act (Wod), by means of review by an external independent Animal Ethics Committee, approval by the Competent Authority and the internal animal welfare body. All animals were housed in Makrolon type III cages, containing Aspen wood shavings, a shelter and nestlets in a controlled environment (12/12 light/dark cycle with lights on at 08:00, 21±2°C) with *ad libitum* access to food (see below) and tap-water. Experimental animals were bred in-house. As the bodyweight, body composition and behavior of mice can vary between sexes, and it was beyond scope to compare male and female groups, only animals of one sex (male offspring) were used as experimental animals. Dams and female offspring were present during breeding and after birth and were removed from the study at weaning age (see below). Primiparous breeder dams and males were obtained from a commercial breeder (Study 1 and 2: C57Bl/6JOlaHsd, Envigo, The Netherlands; Study 3: C57Bl/6J, Charles River Laboratories, Germany). Prior to breeding, females were housed in pairs and males were housed individually to prevent fighting. Breeding was performed by introducing one (Study 3) or two females (Study 1 and 2) into the cage of a male mouse for three consecutive days; all animals were thereafter returned to their home cage. Females were individually housed from 14 days after breeding and were provided with nest material (Nestlet) and left undisturbed until birth of the litter. The day that the litter was born was considered postnatal day (PN) 0.

### Experimental design and outcome parameters

#### Study 1

To induce impaired n-3 accumulation in the offspring brain, females were switched at 4 weeks prior to breeding to a semi-synthetic American Institute of Nutrition-93 Growth (AIN-93G; (36)) based rodent diet with 7% lipids containing 2.58% C18:2n-6 and 0.01% C18:3n-3 as source of n-6 and n-3 PUFAs respectively (n-3 deficient diet; n-3DEF1.). The **n-3DEF1** diet was continued throughout pregnancy and lactation. Male breeders were kept on AIN-93 Maintenance (AIN-93M). At PN2, litters were culled to six pups per dam (each litter containing at least two males and two females). From PN16 onwards, litters (with dam) were randomly assigned to receive either an experimental diet with added Egg-PL as ingredient (ingredient, ING1) or a diet with the same composition, but with an adapted supramolecular lipid structure (structure, STR1). At PN21 the dams and female offspring were removed and the male offspring (ING1, n = 4 litters [12 offspring]; STR1, n = 5 litters [15 offspring]) remained housed with their littermates and their respective diets continued. At PN28 the offspring were housed individually. At P42, animals were euthanized by deep anesthesia (isoflurane N_2_O-O_2_) followed by decapitation. The liver and brain were harvested and the hippocampus was isolated from one hemisphere. All tissues were snap frozen (−80° C). Frozen livers and intact brain hemispheres were homogenized in phosphate buffered saline (PBS), and tissue fatty acid composition was analyzed using gas chromatography, as previously described (37). Tissue fatty acid classes and individual fatty acid species were expressed as a percentage of total fatty acids (% FA). As exploratory readouts, mRNA expression of myelin-associated glycoprotein (MAG), synaptophysin (SYP) and ionized calcium binding adaptor molecule 1 (Iba1) were determined in the hippocampus as molecular markers for myelin integrity, synaptic plasticity and microglia activation respectively; these are also known to be disrupted by n-3 deficiency during development (38–40). Total ribonucleic acid (RNA) from the hippocampus was isolated with the RNeasy mini kit (©QIAGEN) according to the manufacturer’s instructions. Isolated RNA was used to synthesize complementary deoxyribonucleic acid (cDNA) with the iScript^tm^ cDNA Synthesis Kit (Bio-Rad) according to the manufacturer’s instructions. Quantitative polymerase chain reaction (qPCR) was performed with the obtained cDNA, SYBR™ Select Master Mix (Applied Biosystems™), and validated primers to quantify gene expression of MAG (F-CCTGGATCTGGAGGAGGTGA, R-TTCACTGTGGGCTTCCAAGG), IBA1 (F-GATTTGCAGGGAGGAAAAGCT, R-AACCCCAAGTTTCTCCAGCAT), SYP (F-GAGGGACCCTGTGACTTCAG, R-AGCCTGTCTCCTTGAACACG) and housekeeping genes 18s, Gusb, HPRT1, B2M, TBP, Tubb3, and Actin. Gene expression of MAG, IBA1 and SYP was analysed relative to housekeeping genes with the qBase+ software (biogazelle version 3.3). Due to the experimental design in this study involving maternal diet exposure during pregnancy and lactation, the litters rather than individual pups were regarded as experimental units. Tissue fatty acids and gene expression levels were therefore expressed per litter (i.e., average of all individuals within litter). A group of offspring born from n-3DEF1 dams and kept on n-3DEF1 diet until PN42 were included in the study as a deficient reference group (DEF1, n = 2 litters [7 offspring]) for tissue fatty acid composition. In addition, a group of mice born from dams that were kept during (pre-) pregnancy and lactation on an n-3-sufficient control diet (AIN-93G based rodent diet with 7% fat containing 2.63% C18:2n-6 and 0.45% C18:3n-3 as source of n-6 and n-3 PUFAs, respectively), and exposed to a diet containing standard infant formula between PN16 and PN42, was included in the study as a reference for tissue fatty acid composition of (non-deficient) male C57Bl/6J mice (REF1, n = 6 litters [17 offspring]).

#### Study 2

To study effects of early life dietary lipid structure on adult sensitivity to diet-induced obesity, an experimental design was used as previously published (41). In short, breeding females were kept on semi-synthetic AIN-93G diet starting two weeks prior to breeding. At PN2, litters were randomized and culled to 6 pups per dam (2 females, 4 males) to standardize litter size and composition as these factors may affect sensitivity to later in life (diet induced) obesity (42). From PN16 onwards, litters (with dam) were randomly assigned to receive either an experimental diet with added MFGM as ingredient (ING2) or a diet with the same composition and adapted supramolecular lipid structure (STR2). At PN21, male offspring (ING2, n = 12; STR2, n = 12) were weaned and housed in pairs (with littermate) for the remainder of the study. The animals were maintained on their respective experimental diet until PN42 and were thereafter exposed to a semi-synthetic Western style diet (WSD, 20%, w/w fat) until the end of the study at PN98. At PN42, PN70 and PN98, animals were briefly anaesthetized (isoflurane N_2_O-O_2_) to allow body composition analysis using dual-energy X-ray absorptiometry scanning (fat mass and lean mass; DEXA, PIXImus imager, GE Lunar). Relative fat mass was calculated as a percentage of body weight (% fat). A group of mice, raised on a diet containing standard infant formula between PN16 and PN42 and exposed to a standard low fat maintenance diet (AIN93-M) during adulthood, was included in the study as a reference (REF2, n = 12) for normal body composition development of male C57Bl/6J mice on standard (low fat) semi-synthetic diet (41).

#### Study 3

Breeder animals were kept on a standard grain based rodent diet (Altromin 1310M, Germany) during breeding and lactation. At PN2, litter size was standardized to 3 pups per dam (2 males, 1 female or 1 male, 2 females), which is considered a small litter size for C57Bl/6J mice. Small litter rearing in rodents is a paradigm known to induce predisposition towards obesity (42). From PN16 onwards litters (with dam) were randomly assigned to receive an experimental diet with added MFGM as ingredient (ING3) or a diet with the same composition and adapted supramolecular lipid structure (STR3). At PN21 male offspring were weaned and housed in pairs (with littermate) for the remainder of the study (ING3, n = 10; STR3, n = 12). Animals were kept on their respective experimental diet until PN42 and were thereafter exposed to AIN-93M diet until PN98, the end of the study. At PN42 and PN98 the body composition of the mice was determined by EchoMRI Whole Body Composition Analyser (EchoMRI, Houston, FL, USA). Relative fat mass was calculated as a percentage of body weight (% fat). At PN70 and PN71 animals were subjected during the light phase to an open field test (OF) to assess exploratory activity and a novel object recognition test (NOR) to assess memory performance as previously described (see (43)) with some adaptations to the protocols. Briefly, the OF test comprised a five-minute trial in which mice were introduced to the center of a square test area (50 x 50 cm) and allowed to explore the area. The total distance moved, and the time spent in the center square of the arena, were recorded using Ethovision^®^ XT 11.5 (Noldus, Wageningen, The Netherlands), using the nose, center and tail base of the mouse for tracking. Prior to each test, mice were housed individually for 30 minutes, in a clean cage in the room where behavior testing took place. Cage mates were tested consecutively, but cages with mice were taken at random from the animal room. Mice were returned to their home cage after the tests. During the first trial of the NOR, which started 60 min after the OF, mice were introduced again to the same test area in which there were now two identical objects, placed in opposite corners of the area. Mice were allowed to explore the area and the objects for five minutes and were returned to their home cage thereafter. On the next day (test trial), this procedure was repeated, but one of the two objects in the test area was replaced by a new object which was different in shape but similar in size and material. The time that was spent exploring each object during the test trial was recorded. Performance was evaluated by calculating a recognition index: (*N/(N+F*)) where *N* is time spent exploring the novel object and *F* is time spent exploring the familiar object. A higher index reflects better memory performance. A group of mice that were raised in a normal size litter before weaning (n = 6 pups per litter and litters containing at least 2 males and 2 females) and exposed to a diet containing standard infant formula between PN16 and PN42, followed by AIN-93M during adulthood, was included in the study as a reference (REF3, n = 12) for normal body composition development of male C57Bl/6J mice.

### Experimental Diets

All diets used in Study 1 were supplied by Sniff Spezialdiäten (Soest, Germany), whilst diets used in Studies 2 and 3 were supplied by Research Diet Services (Wijk bij Duurstede, the Netherlands), unless specified otherwise. The experimental diets were provided to the animals in the form of a soft dough ball that was freshly prepared on daily basis and placed on the cage floor to allow easy access for the animals. The ING and STR experimental diets were semi-synthetic with a macronutrient composition according to AIN-93G as previously described (37, 41, 44). In short, the experimental diets consisted of 28.3% w/w IF powder complemented with additional protein, carbohydrates and micronutrients to mimic AIN-93G composition. The lipid fraction in the experimental diets was entirely derived from the IF powders. The IF powders contained phospholipids (PL, 0.4 g/100g powder) either sourced from egg yolk (Study 1) or MFGM-rich beta-serum and were produced in a pilot plant (Danone Nutricia Research, Utrecht, The Netherlands) using the same recipe and ingredients per study. ING and STR experimental diets were therefore always equal in nutrient composition but differed from each other *only* in the supramolecular structure of lipid droplets as a result of an adapted manufacturing process for the IF powder, as previously described (44, 45). Lipid droplets in STR diets were large in size, with thin interfaces likely covered by phospholipids, while ING diets comprised small lipid droplets (Table 1); although phospholipids were present, the interface of the lipid droplets was composed of proteins (Fig 2).

**Table 1.**
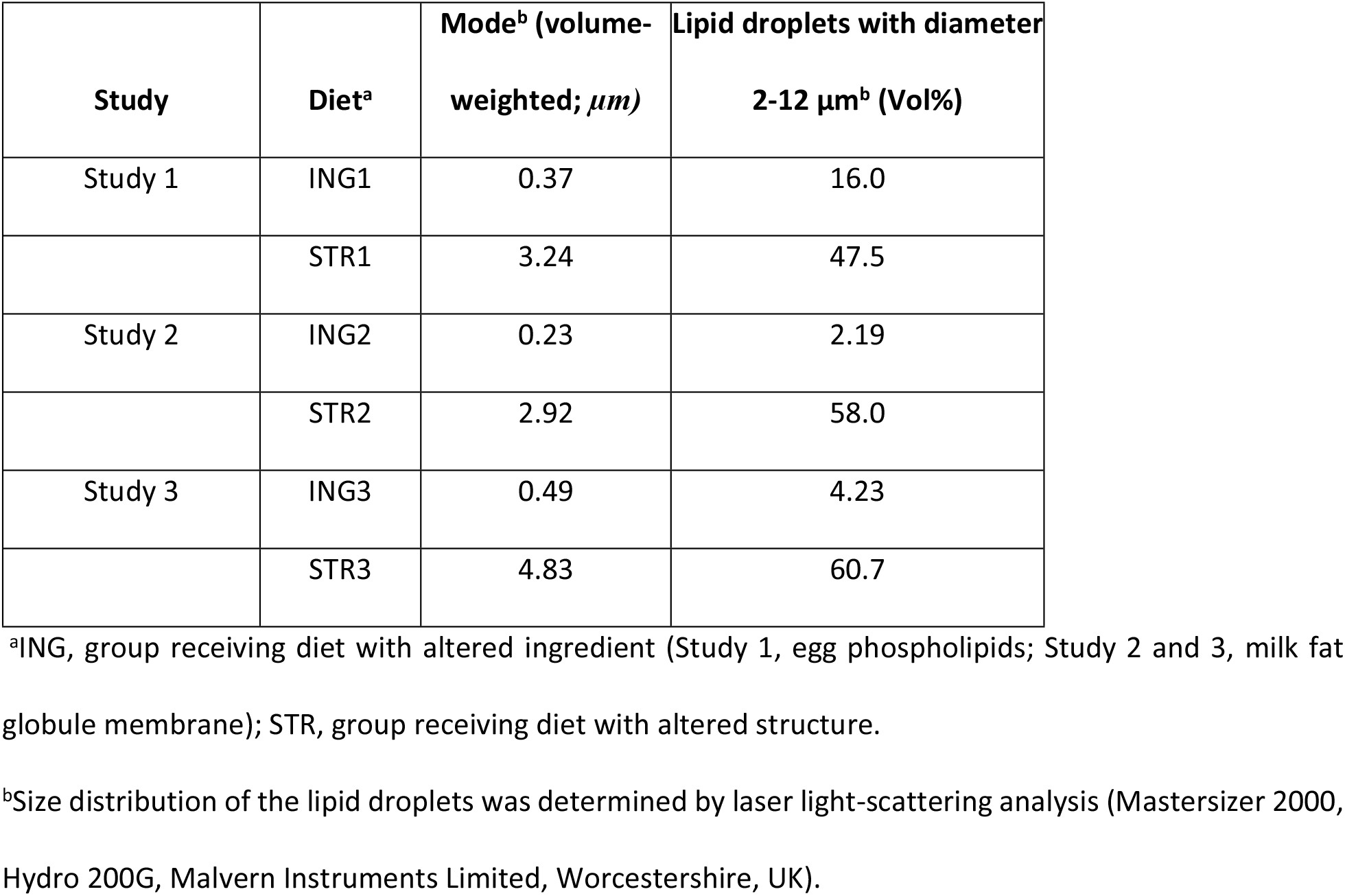
Particle size distribution of lipid droplets in experimental diets.

**Figure 2.**
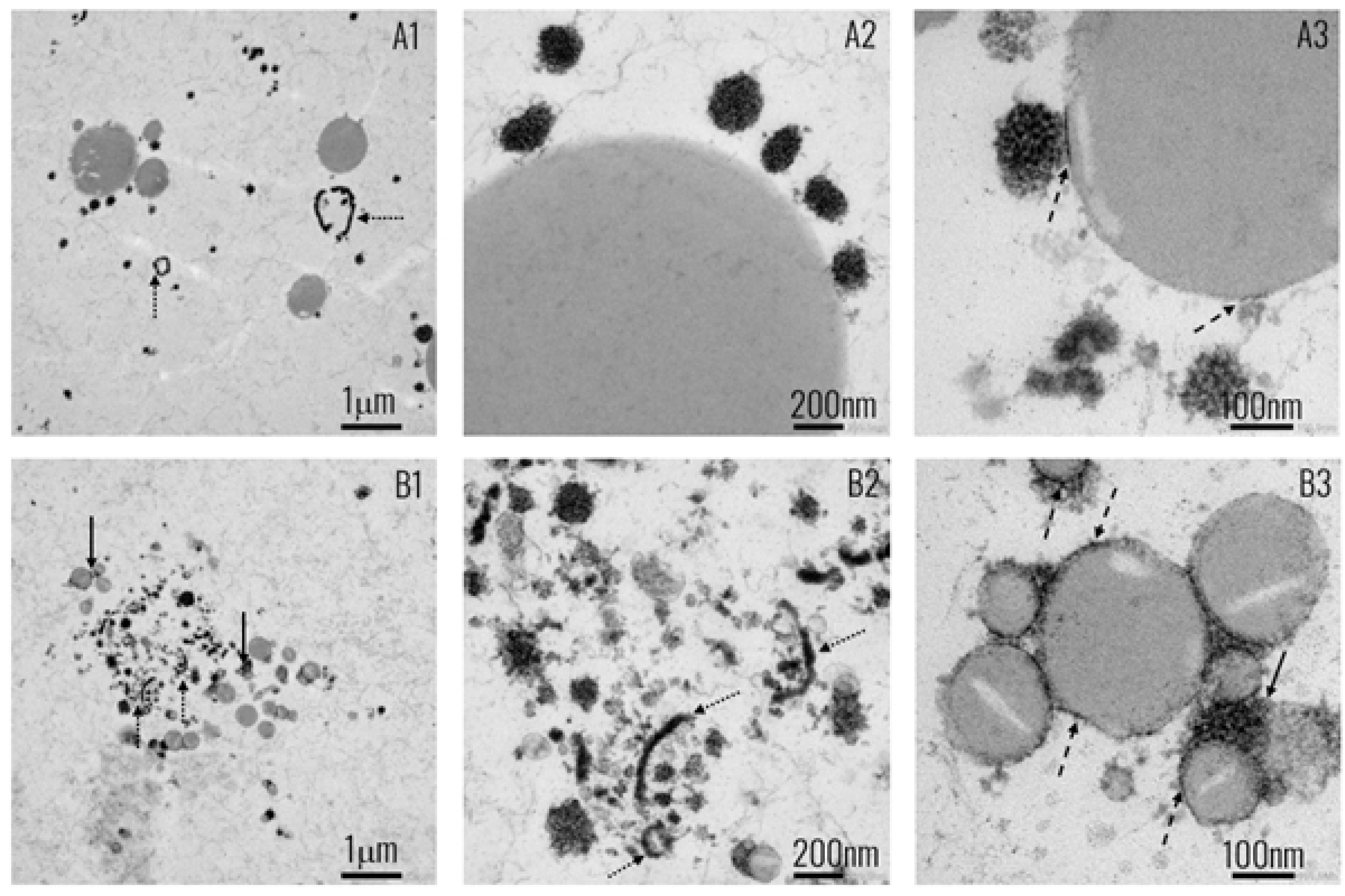
Transmission electron microscopy (TEM) images of experimental diets STR (A) and ING (B). A1-3 Large sized fat droplets displaying a thin interface mainly, indicating the presence of phospholipids. A2-3 a close-up on the interface of a lipid droplet with few associated casein micelles. B1 Small sized lipid droplets aggregated by proteins with MFGM fragments in the aqueous phase (B2). B3 A close-up on clustering of lipid droplets and their interfaces composed of protein. Solid arrows are pointing at large casein/protein aggregates, clustering lipid droplets; the dashed arrows are pointing at interfacial protein; the dotted arrows are pointing at MFGM fragments or vesicles in the aqueous phase. Sample preparation and TEM imaging was performed as previously described (45).

### Sample size and statistical analysis

number of litters that were generated after breeding. To determine the sample sizes required for the experimental groups in Study 2 and 3, power calculations were performed based on published data from previous experiments with comparable design and diet interventions (44, 46). Using an error-probability of 5% and power of 80% the sample sizes were calculated to be 12 animals per group for both studies. In study 3, one animal in the ING3 group was taken out of the experiment shortly after weaning due to malocclusion, its cage mate was removed as well, resulting in a group size of 10. Data were analyzed by an experimenter blinded to the group allocation. All data were analyzed with SPSS 19.0 software (SPSS Benelux, Gorinchem, The Netherlands). In Study 1, data were analyzed using one way ANOVA with litters regarded as experimental unit rather than individual animals due to the maternal exposure to n-3 deficient diet. In Study 2 and 3, individual animals were considered as experimental units and data were analyzed using linear mixed models that included litter of origin and/or home cage i.e., the shared environment between individuals within groups, as random factor in the model. Fixed factors for all studies included group, and for Study 2 additional fixed factors were time (repeated) and its interaction with experimental diet. When applicable, significant main effects of group and/or group interaction effects were followed by post-hoc comparisons (Least Squares Difference, LSD) between experimental groups. Direct comparisons between experimental diet groups and reference groups were omitted as these comparisons were not meaningful due to different nutrient composition of the diets. All data are presented as mean ± SEM, differences between ING and STR were considered significant at p < 0.05.

## RESULTS

### Study 1

At P42, groups did not differ in body weight (g) (n-3DEF1, 23.2 ± 0.27; ING1, 22.0 ± 1.67; STR1, 22.3 ± 0.45; REF1, 22.9 ± 0.51). Liver and brain tissue fatty acid composition are presented in Table 2 and 3 respectively. Exposure to n-3DEF1 diet resulted in lower tissue levels of linoleic acid (LA) and n-3 long chain polyunsaturated [(LC)PUFA] species, and increased in n-6 docosapentaenoic acid (DPA) and other n-6 PUFA species in brain specifically. Animals that were kept on STR1 compared to ING1 diet after maternal alpha-linolenic (ALA) deficiency showed significantly higher levels of LA and the n-3 PUFA species ALA, eicosapentaenoic acid (EPA) and n-3 DPA in liver tissue (Table 2). In brain, STR1 resulted in higher total n-3 (LC)PUFA, including DHA, and lower total n-6 (LC)PUFA levels, including arachidonic acid and n-6 DPA (Table 3). mRNA expression of myelin-associated glycoprotein (MAG) in the hippocampus was increased in animals on STR1 compared to ING1 (F_(1,7)_=6.74, p = 0.04, Fig 3A), but hippocampal expression of synaptophysin (SYP) and ionized calcium-binding adaptor molecule 1 (Iba 1) remained unaffected by diet (Fig 3B and C).

**Table 2.** Fatty acid composition in liver.

| Liver fatty acids |  | n-3DEF1 | ING1 | STR1 | REF1 |
| --- | --- | --- | --- | --- | --- |
| SFA | | $25.7 \pm 1.07^a$ | $31.8 \pm 1.01$ | $32.5 \pm 0.67$ | $33.3 \pm 1.41$ |
| MUFA | | $57.2 \pm 2.43$ | $47.4 \pm 2.72$ | $42.4 \pm 2.19$ | $40.2 \pm 4.23$ |
| PUFA | | $17.0 \pm 1.34$ | $20.7 \pm 1.75$ | $25.0 \pm 1.91$ | $26.4 \pm 2.91$ |
| N-6 | C18:2 n-6 | $5.24 \pm 0.03^a$ | $5.47 \pm 0.44^b$ | $6.96 \pm 0.42$ | $7.01 \pm 0.48$ |
| | C20:4 n-6 | $7.28 \pm 1.09$ | $7.78 \pm 0.72$ | $8.27 \pm 0.76$ | $8.75 \pm 1.13$ |
| | C22:4 n-6 | $0.24 \pm 0.01$ | $0.23 \pm 0.01$ | $0.24 \pm 0.02$ | $0.28 \pm 0.02$ |
| | C22:5 n-6 | $1.44 \pm 0.03^a$ | $0.41 \pm 0.04$ | $0.31 \pm 0.03$ | $0.35 \pm 0.05$ |
| | Sum n-6 | $15.9 \pm 1.11$ | $15.4 \pm 1.25$ | $17.3 \pm 1.24$ | $17.9 \pm 1.60$ |
|  | **Sum n-6**<br>**LCPUFA** | 10.5 ± 1.18 | 9.77 ± 0.81 | 10.1 ± 0.85 | 10.7 ± 1.13 |
| **N-3** | **C18:3 n-3** | 0.09 ± 0.01<sup>a</sup> | 0.21 ± 0.01<sup>b</sup> | 0.40 ± 0.02 | 0.35 ± 0.02 |
|  | **C20:5 n-3** | 0.00 ± 0.00<sup>a</sup> | 0.21 ± 0.03<sup>b</sup> | 0.48 ± 0.06 | 0.41 ± 0.09 |
|  | **C22:5 n-3** | 0.02 ± 0.01<sup>a</sup> | 0.13 ± 0.01<sup>b</sup> | 0.24 ± 0.02 | 0.26 ± 0.05 |
|  | **C22:6 n-3** | 0.89 ± 0.23<sup>a</sup> | 4.70 ± 0.47<sup>b</sup> | 6.47 ± 0.62 | 7.33 ± 1.18 |
|  | **Sum n-3** | 1.03 ± 0.23<sup>a</sup> | 5.29 ± 0.51<sup>b</sup> | 7.66 ± 0.68 | 8.41 ± 1.32 |
|  | **Sum n-3**<br>**LCPUFA** | 0.92 ± 0.24<sup>a</sup> | 5.07 ± 0.51<sup>b</sup> | 7.24 ± 0.66 | 8.05 ± 1.31 |

**Table 3.**
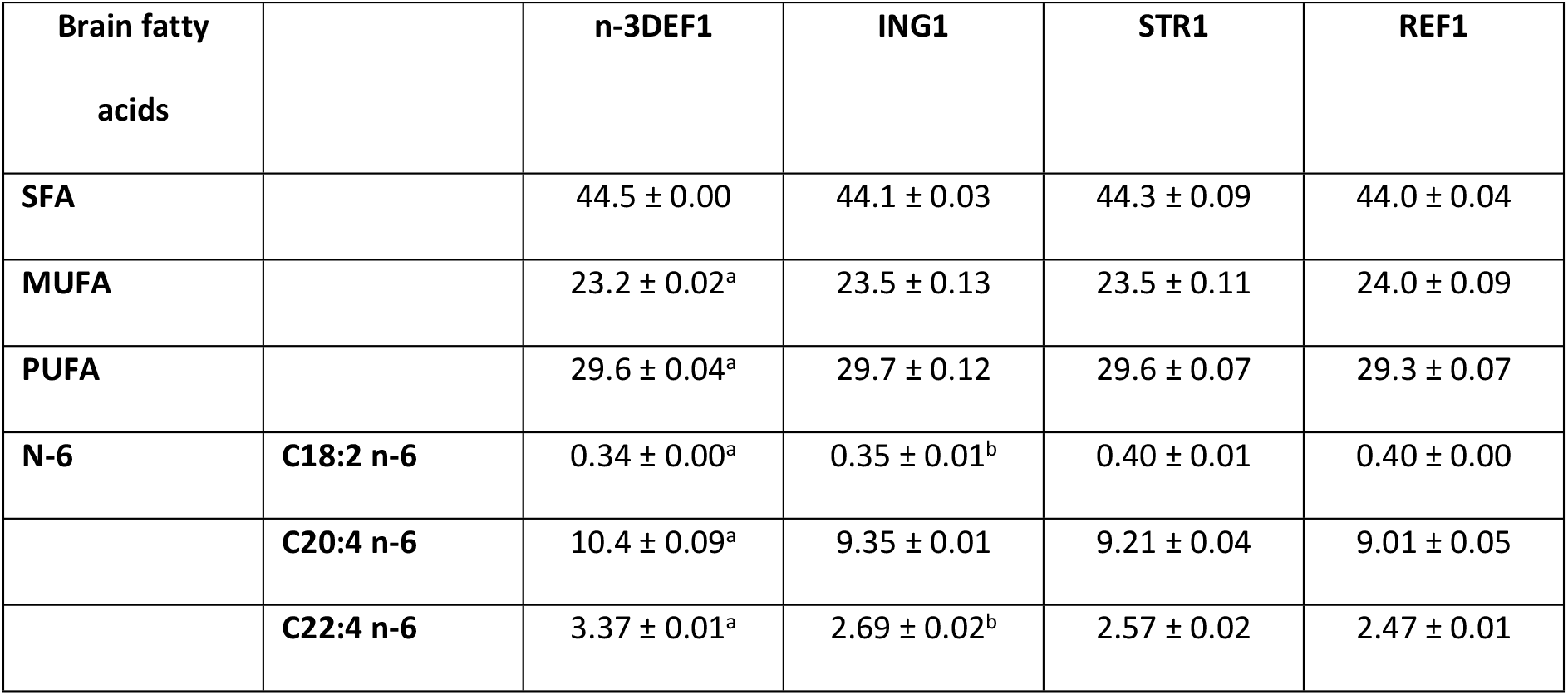
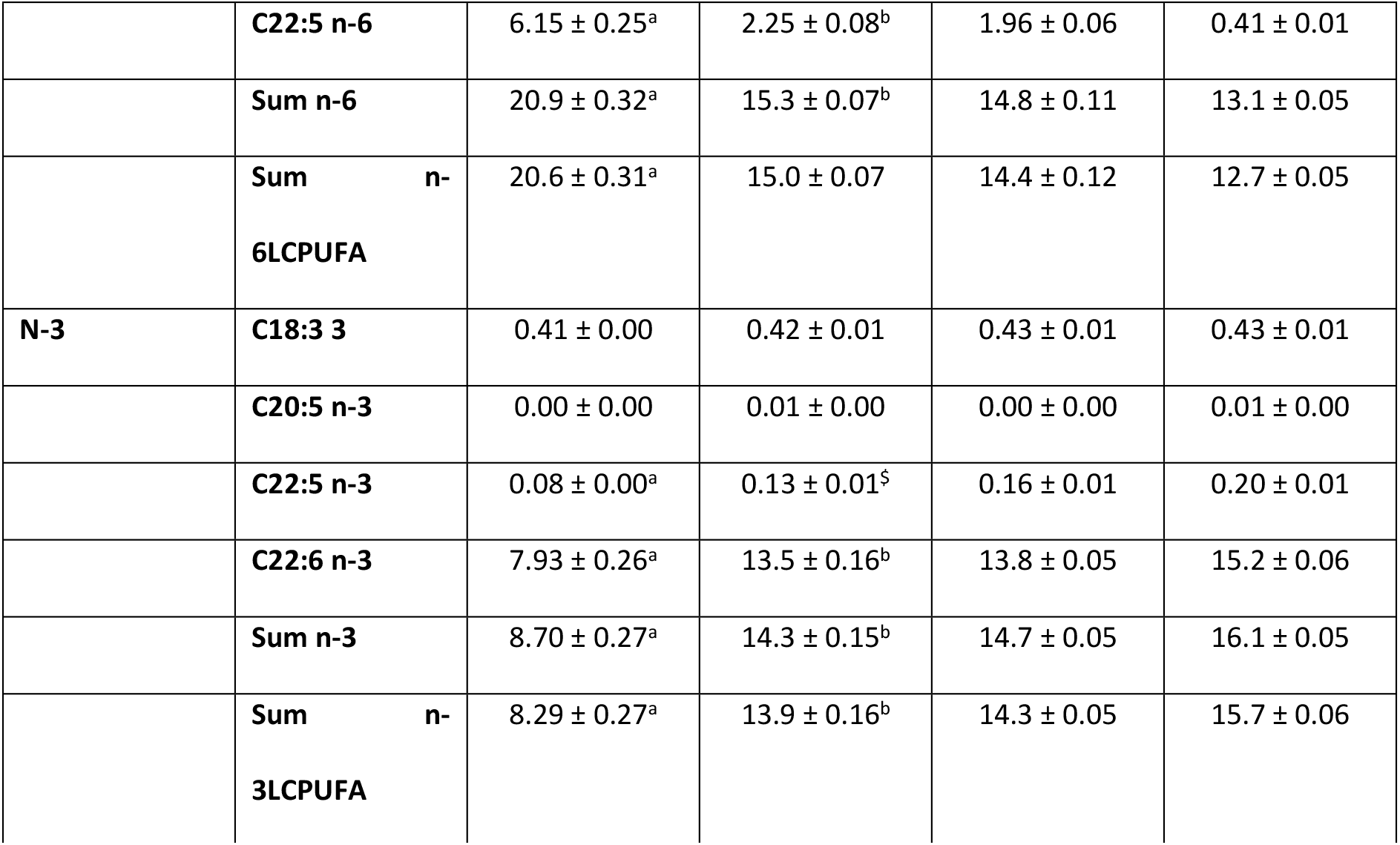
Fatty acid composition in brain tissue.

**Figure 3.**
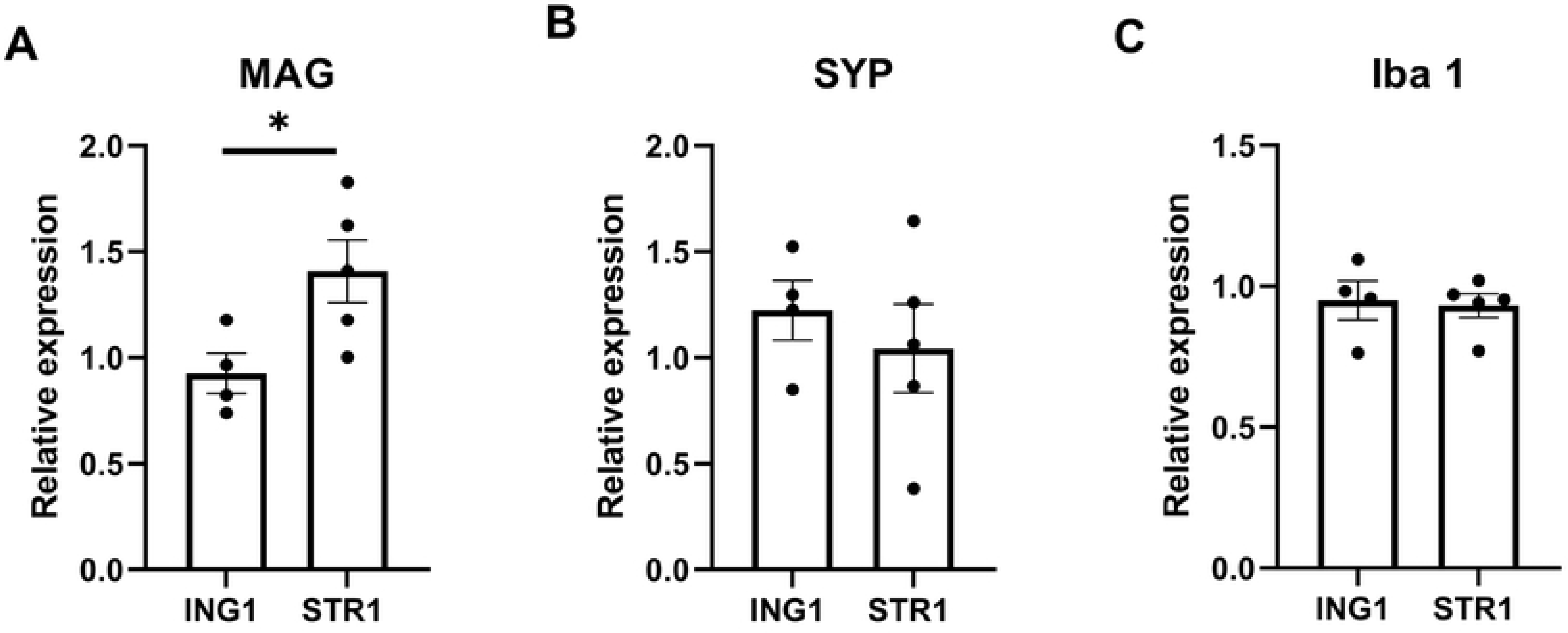
Relative mRNA expression of MAG, SYP and Iba 1 in hippocampus of mice exposed to experimental diets following previous maternal n-3 deficiency. Relative mRNA expression in hippocampus of mice exposed to experimental diet (P16-42) with added phospholipid ingredient (ING, n = 4 litters [12 mice]) and experimental diet with adapted supramolecular lipid structure (STR1, n = 5 litters [15 mice]). MAG, myelin-associated glycoprotein, Syp, Synaptophysin; Iba 1, ionized calcium-binding adaptor molecule 1; ING1, group receiving diet with altered ingredient (Study 1, egg phospholipids); STR, group receiving diet with altered structure. Data are means ± SEM; ^*^ significant difference between ING and STR group (p < 0.05)

Fatty acids (% of total fat) in liver of mice exposed to n-3 deficient diet (n-3DEF1, n = 2 litters [7 mice]), experimental diet (P16-42) with added phospholipid ingredient (ING1, n = 4 litters [12 mice]) and experimental diet with adapted supramolecular lipid structure (STR1, n = 5 litters [15 mice]) or healthy reference diet (REF1, n = 6 litters [19 mice]). FA, fatty acids; (LC)PUFA, (long chain) poly unsaturated fatty acids; MUFA, monounsaturated fatty acids; SFA, saturated fatty acids; ING1, group receiving diet with altered ingredient (Study 1, egg phospholipids); STR1, group receiving diet with altered structure; REF1, reference diet. Data are means ± SEM. ^a^ significant difference between n-3DEF1 and REF1 groups (p < 0.05); ^b^ significant difference between ING1 and STR1 groups (p < 0.05).

Fatty acid (% of total fat) in brain tissue of mice exposed to n-3 deficient diet (n-3DEF1, n = 2 litters [7 mice]), experimental diet (P16-42) with added phospholipid ingredient (ING1, n = 4 litters [12 mice]) and experimental diet with adapted supramolecular lipid structure (STR1, n = 5 litters [14 mice]) or healthy reference diet (REF1, n = 6 litters [18 mice]). FA, fatty acids; (LC)PUFA, (long chain) polyunsaturated fatty acids; MUFA, monounsaturated fatty acids; SFA, saturated fatty acids; ING, group receiving diet with altered ingredient (Study 1, egg phospholipids); STR1, group receiving diet with altered structure; REF1, reference diet. Data are means ± SEM. ^a^ significant difference between n-3DEF1 and REF1 groups (p < 0.05); ^b^ significant difference between ING1 and STR1 groups (p < 0.05); ^$^trend towards difference between ING1 and STR1 groups (0.05 < p ≤ 0.06).

### Study 2

At P42, the final day of the early life diet intervention, there were no differences between groups exposed to ING2 and STR2 in body weight (g) and body composition. However, body weight gain on WSD diet during adulthood (from P42 to P98) was lower in the group that had previously been exposed to STR2 compared to ING2 (diet*week interaction (F_(4,33)_=15.689, p < 0.01), Fig 4A). DEXA scans revealed that, while lean mass development over time was not significantly different (Fig 4B), animals that were raised on STR2 showed reduced adult body fat accumulation compared to animals that had been raised on ING2 (diet*week interaction: fat mass, [F_(4,33)_=12.455, p < 0.01]; relative fat mass, [F_(4,33)_=8.483, p < 0.01], Fig 4C and D).

**Figure 4.**
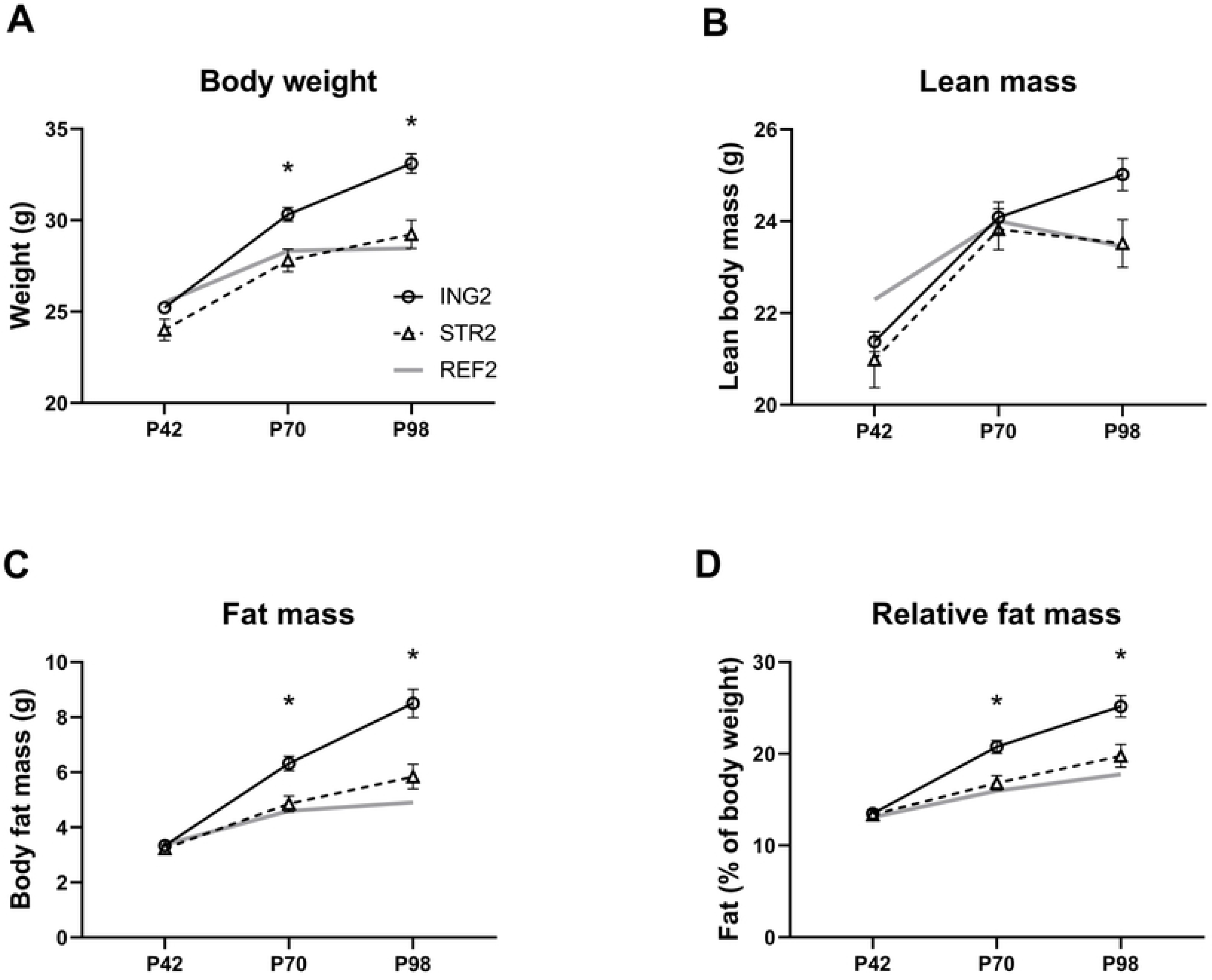
Body weight and body composition development during WSD challenge (week 6-14) of mice that had previously been exposed to experimental diets. (A) body weight, (B) lean mass, (C) fat mass and (D) relative fat mass of adult mice on Western style diet (weeks 6 - 14) that were previously exposed (postnatal day 16-42) to experimental diet with added phospholipid ingredient (ING2, n = 12) or experimental diet with adapted supramolecular lipid structure (STR2, n = 12). A group of mice that were raised on a neutral diet and exposed to low fat AIN-93-M were monitored and included as a reference for body weight and body composition development under non-challenged conditions (REF2, n = 12). ING2, group receiving diet with altered ingredient (Study 2, milk fat globule membrane); STR2, group receiving diet with altered structure; REF2, reference diet; P, postnatal day. Data are means ± SEM; ^*^significant difference between ING and STR group (p < 0.05).

### Study 3

At P42, there were no differences between the ING3 and STR3 exposed groups in body weight or body composition (Fig 5A–D). While body weight and lean body mass were not different between groups in adulthood at PN98 and the difference between groups on fat mass did not reach significance (Fig 5A, B and C respectively), the relative fat mass of STR3 exposed animals at P98 was lower compared to that of ING3 ([F_(1, 9)_=6.869, p = 0.028], Fig 5D). During the open field test at PN 70 body weight was not different between diet groups (ING3, 27.1 ± 0.76 g; STR 28.6 ± 0.31 g) and groups did not display differences in distance travelled or time (%) spent in the center square (Figs 6A and B). The recognition index during the novel object recognition test at P71 was, however, significantly higher in animals previously exposed to STR3 compared to ING3 (F_(2,31)_=3.364, p = 0.48, Fig 6C).

**Figure 5.**
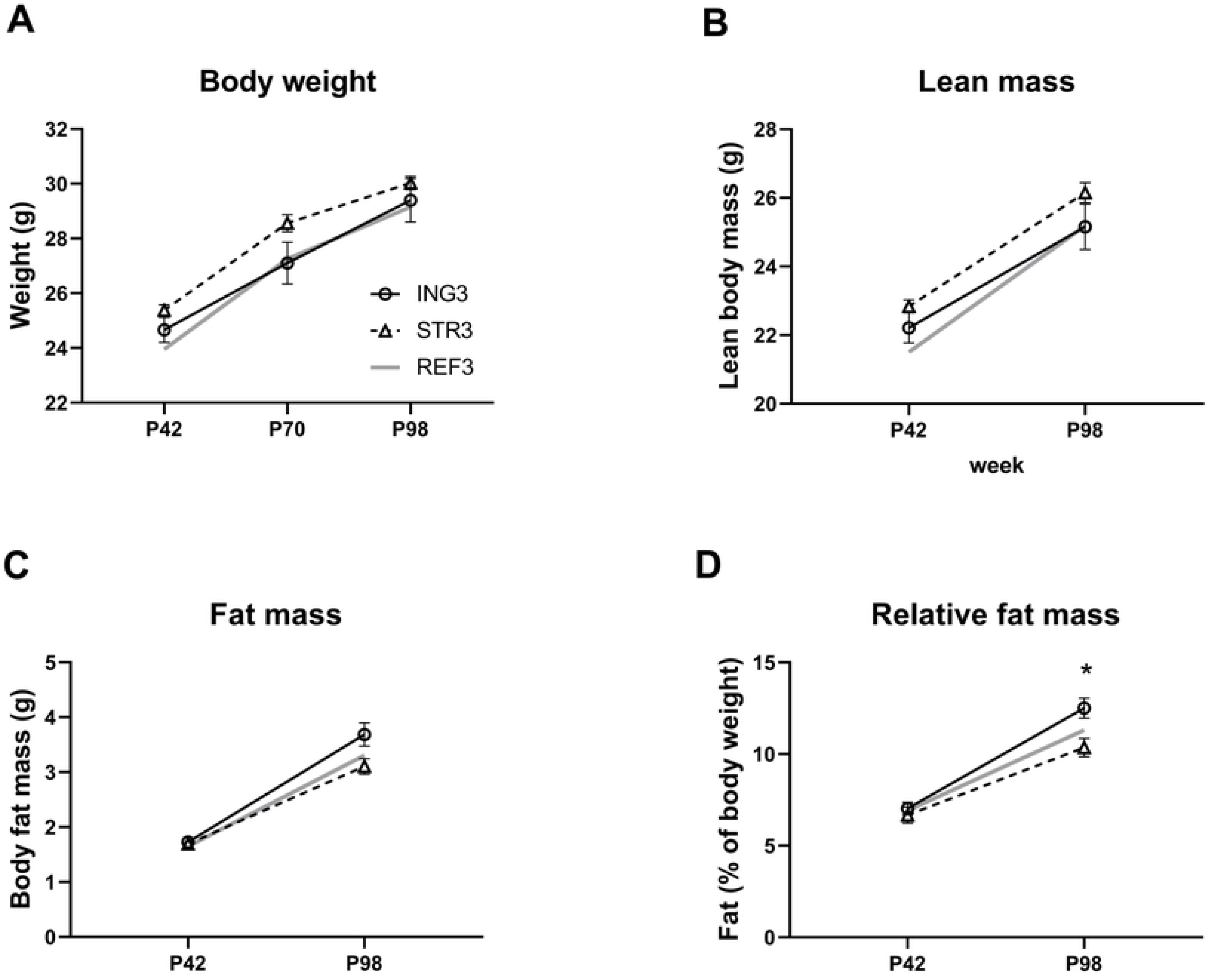
Body weight and body composition at day 98. (A) body weight, (B) lean body mass (C) fat mass (D) relative fat mass of adult male mice that had previously been exposed (postnatal day P16-42) to experimental diet with added ingredient (ING3, n = 10) or experimental diet with adapted structure (STR3, n = 12). ING3, group receiving diet with altered ingredient (Study 3, milk fat globule membrane); STR3, group receiving diet with altered structure; REF3, reference diet; P, postnatal day. Data are means ± SEM; ^*^ = significant difference between ING3 and STR3 (p < 0.05).

**Figure 6.**
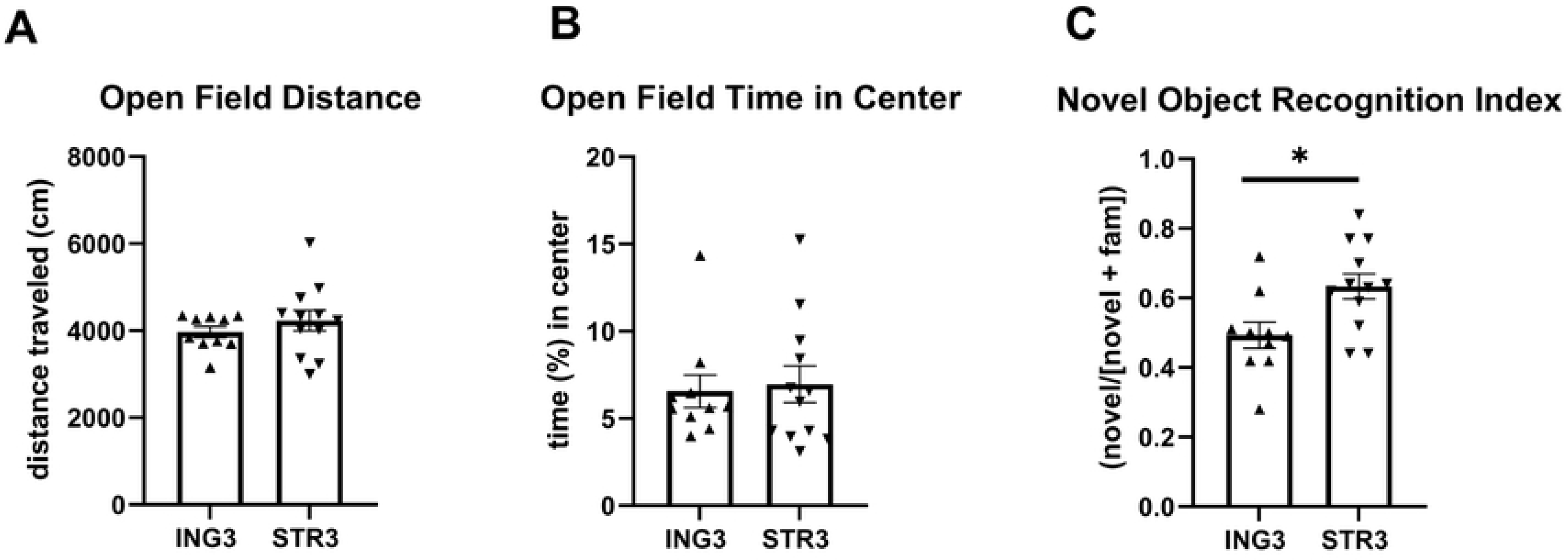
Exploration behavior in the open field test and memory performance in novel object recognition test. (A) total distance moved in open field, (B) relative time spent in center of the open field and (C) novel object recognition index of adult that had previously been exposed (postnatal day 16-42) to experimental diet with added ingredient (ING3, n = 10) or experimental diet with adapted structure (STR3, n = 12). ING3, group receiving diet with altered ingredient (Study 3, milk fat globule membrane); STR3, group receiving diet with altered structure. Data are means ± SEM; ^*^ = significant difference between ING3 and STR3 (p < 0.05).

## Discussion

Here, we have demonstrated for the first time that the supramolecular structure of lipids in the early life diet has the capacity to impact various (long-term) health outcomes. Mice that were exposed in early life to IF in which lipids were organized as large, phospholipid-coated lipid droplets showed better incorporation of n-3 (LC)PUFA in liver and brain tissue during development, increased hippocampal expression of MAG, reduced adult body fat accumulation in different obesity models and improved adult cognitive function, compared to mice that had been exposed to IF in which the *exact same ingredients* were present, but that lacked the complex supramolecular lipid structure. In other words, when the composition, size *and* structure of the lipids was closer to the structure of human MFG, health benefits resulted. These data suggest that the typical supramolecular structure of lipid globules themselves may be a significant factor contributing to the lifelong advantages observed in BF compared to FF infants.

In previous studies, using similar programming models, we have comprehensively shown that early life exposure to concept IFs containing large, MFGM coated lipid droplets resulted in beneficial effects on later life body composition (41, 44, 47, 48), metabolic health (49–54) and brain outcomes (43, 46, 55, 56), when compared to IFs with small lipid droplets and no MFGM added. However, due to the experimental design of these studies, the effects of the mere presence of MFGM versus effects of the complex supramolecular lipid droplet structure with large lipid droplets and MFGM fragments at the interface could not be distinguished. The results from the current experiments suggest that these previous findings were, in part, mediated by the modified dietary lipid structure and that we should not underestimate the role that the supramolecular lipid droplet structure of IF plays in programming later life health.

Clinical studies with IFs supplemented with MFGM have shown promising effects on brain and immune outcomes, yet the evidence base is still small and heterogeneous due to differences in: MGFM sources, formula composition, intervention duration and methodologies used to measure outcomes at different timepoints (28). Some studies have shown effects of supplementation with MGFM, sometimes with other added bioactive ingredients, on neurocognitive and language scores at 6 (57) and 12 months (58–60), and at 4 years of age (61). Although there are some ambiguous outcomes, there is also clinical evidence on reductions in otitis media, upper respiratory tract infections and antipyretic use (28). The clinical growth equivalence and tolerance studies published to date show that MGFM, as an ingredient, can be safely added to IF, yet there is no evidence that the mere presence of MFGM in IF modulates infant growth trajectories or has a protective effect against overweight or metabolic disturbances. Several studies show similar growth patterns, both weight and height, at 4, 6, 12 and 18 months (58–60, 62–64). In addition, some studies show differences in plasma cholesterol and plasma and erythrocyte polar lipids between experimental and control IF, but these differences were no longer observed at 12 months (65). The current experiments suggest that, aside from the beneficial impact of MFGM as an ingredient on cognitive and immune outcomes, the specific supramolecular lipid structure in IF with large, MFGM-coated lipid droplets contributes to the wealth of neurocognitive, body composition and metabolic health benefits previously reported (44, 46, 47, 51, 52, 54, 56).

The precise mechanisms by which the supramolecular structure of dietary lipid droplets influences (later life) body composition, metabolic health and neurocognitive function remain to be elucidated, but there are indications from several studies that lipid structure impacts absorption and digestion kinetics, resulting in a different appearance of lipids and (postprandial) hormones in the bloodstream after ingestion (33, 66, 67). Such different bioavailability of lipids may alter their metabolic fate; this is hypothesized to program adipocyte energy metabolism towards reduced lipid storage in white adipose tissue (68). Moreover, any differences in bioavailability of lipids and circulating hormones to the developing brain is hypothesized to affect brain development and neurocognitive function.

Circulating lipids, including (n-3) LCPUFAs, are incorporated into neuronal tissue early in life and drive key neurodevelopmental processes, such as myelination, and hormones derived from other organs and tissues may be used as neurodevelopmental signals (69). Thus, early exposure to a more human-like MFG structure may support brain development and cognitive function via bioavailability of lipids, and program metabolic homeostasis via alterations in energy storage and expenditure.

Across three separate but linked studies, we demonstrate that early life exposure to large lipid droplets with a complex membrane at the interface, which closely mimics the human MFG structure, programs adult brain and metabolic health outcomes in pre-clinical animal models. These effects were *not* generated through the addition of MFGM as an ingredient nor via other differences in dietary composition. Moving beyond the addition of MFGM as an ingredient in IF to a more nuanced physical alteration in the supramolecular structure of lipid droplets in IF offers new opportunities for nutrition during infancy to program later life health.

